# Membrane barrels are taller, fatter, inside-out soluble barrels

**DOI:** 10.1101/2021.01.30.428970

**Authors:** Rik Dhar, Ryan Feehan, Joanna S.G. Slusky

**Affiliations:** Department of Molecular Biosciences, The University of Kansas, 1200 Sunnyside Ave. Lawrence KS 66045-3101; Center for Computational Biology, The University of Kansas, 2030 Becker Dr., Lawrence, KS 66045-7534

## Abstract

Up-and-down β-barrel topology exists in both the membrane and soluble environment. However, β-barrels are virtually the only topology that exist in the outer membrane. By comparing features of these structurally similar proteins, we can determine what features are particular to the environment rather than the fold. Here we compare structures of membrane β-barrels to soluble β-barrels and evaluate their relative size, shape, amino acid composition, hydrophobicity, and periodicity. We find that membrane β-barrels are generally larger than soluble β-barrels in with more strands per barrel and more amino acids per strand, making them wider and taller. We also find that membrane β-barrels are inside-out soluble β-barrels. The inward region of membrane β-barrels have similar hydrophobicity to the outward region of soluble β-barrels, and the outward region of membrane β-barrels has similar hydrophobicity to the inward region of the soluble β-barrels. Moreover, even though both types of β-barrel have been assumed to have strands with amino acids that alternate in direction and hydrophobicity, we find that the membrane β-barrels have more regular alternation than soluble β-barrels. These features give insight into how membrane barrels maintain their fold and function in the membrane.

## Introduction

Membrane β-barrels are found on the periphery of Gram-negative bacteria, mitochondria, and chloroplasts. Their major functions are small molecule transport, structural support, surface binding and enzymatic catalysis.^1^ Although almost all outer membrane proteins are β-barrels, there are multiple classes among these barrels including some classes of barrels that convergently evolved the barrel structure.^2 3^

In order to determine what amino acid features lead to the membrane β-barrel fold, many features of β-barrels have been determined statistically. Membrane β-barrels tend to have hydrophobic and aromatic amino acids on their exterior surface and neutral and hydrophilic amino acids on their interior.^4^ Membrane barrels are known to have a belt of charged amino acids in the phosphate region of their lipopolysaccharide milieu^5^ and have aromatic amino acids that form an aromatic girdle at both sides of the membrane in the headgroup region.^6^ Aromatic amino acids in membrane barrels are often found to be in neighboring strands to glycine residues^7^, likely to facilitate aromatic rescue.^8^

There is a history of understanding the architecture and amino acid composition of helical membrane proteins in comparison to their soluble counterparts.^9^ Doing so allows for the decoupling of amino acid features required for the secondary structure from features required for the protein insertion and stability in the membrane. Here we aim to study membrane β-barrels in comparison with soluble β-barrels of similar topology. By controlling for the topology we can better understand what features of membrane β-barrels are required for maintaining their fold in the membrane.

There are a considerable number of soluble closed up-and-down β-barrel proteins with which to compare their membrane counterparts. The functions of most of these β-barrels can be classified into four different types: green fluorescence protein that consist of a fluorophore prosthetic group attached at the core of a barrel structure^10^, lipocalins and avidins which bind hydrophobic small molecules^11^, and allene oxide cyclase which catalyzes isomerization of fatty acids.^12^

The structure of membrane and soluble β-barrels have been compared before, revealing that the eight stranded membrane barrels have a higher glycine content and are less twisted, more bent, and have smaller pores than their eight stranded soluble β-barrels counterparts.^13^ Also the geometry of the β-barrel differs, especially with respect to their dihedral angles between membrane and soluble β-barrel.^14^ These studies used smaller datasets that also included non-closed soluble barrels.

In this study, we compare a larger set of structurally solved membrane β-barrels to that of comparably folded soluble β-barrels. We investigate the differences in their size, shape, and amino acid composition. We observe that membrane β-barrels tend to be generally larger in strand number and length than soluble β-barrels. While the membrane β-barrels have more small amino acids, the soluble β-barrels have more charged amino acids. Studying their hydrophobicity, we note that membrane β-barrels are inside-out soluble β-barrels. Finally, analyzing the amino acid alternation patterns we find that the membrane β-barrels alternate more than the soluble β-barrels.

## Methods

### Protein Dataset

We created a dataset of comparable membrane and soluble closed β-barrels. To compile our soluble β-barrel dataset (Figure S1), Protein Data Bank (PDB) structures were selected from the SCOPe database.^15^ We searched for folds using the term ‘barrel’ in either the descriptions or comments. 159 folds were returned. The first PDB entry from each of these 159 folds was then manually inspected for closed β-barrel structure. Eight folds were identified that had soluble β-barrel folds with total of 1444 unique PDB IDs. Highly similar PDB structures were identified using Clustal Omega at 95% identity cutoff^16^, and they were removed with preference for keeping un-mutated sequences in the dataset. The mutant variant PDB structures were eliminated by searching their SEQADV records for following comments: “ENGINEERED”, “ENGINEEREDMUTATION”, “DELETION”, “INSERTION”, or “CONFLICT”. The resulting 273 PDB structures were then manually checked to remove any non-barrel structures. The remaining 269 PDB structures were further refined by removing similar proteins by using PISCES at 50% identity cutoff^17^, resulting in a final set of 101 soluble β-barrel structures (Table ST1). Our membrane dataset is composed of 110 monomeric and 22 polymeric β-barrel with maximum of 50% identity among them (Table ST2) and was collected as previously described.^3a^

### Structure analysis

To identify the barrel regions of each protein, our in house software, Polar Bearal^5 3a^, was used (Figure 1A). The β-strands of the β-barrel in each PDB file were identified and the residues within these β-strands were classified to be inwards or outwards based on their direction relative to axis of the β-barrel. For each β-barrel, strand number, strand length, and mean tilt angles were calculated. As Polar Bearal was optimized to process membrane β-barrels, it needed to be updated to process soluble β-barrels. Processing soluble β-barrels differed in two ways. First, membrane β-barrels are more structurally similar to each other: sequential β-strands are contiguous, and the strand length and shape of membrane β-barrels are generally uniform, likely due to membrane constraints. In contrast, soluble β-barrel strands are not always sequentially contiguous nor are the barrels as uniform, causing misidentification of residue direction in soluble β-barrels.

**Figure 1.**
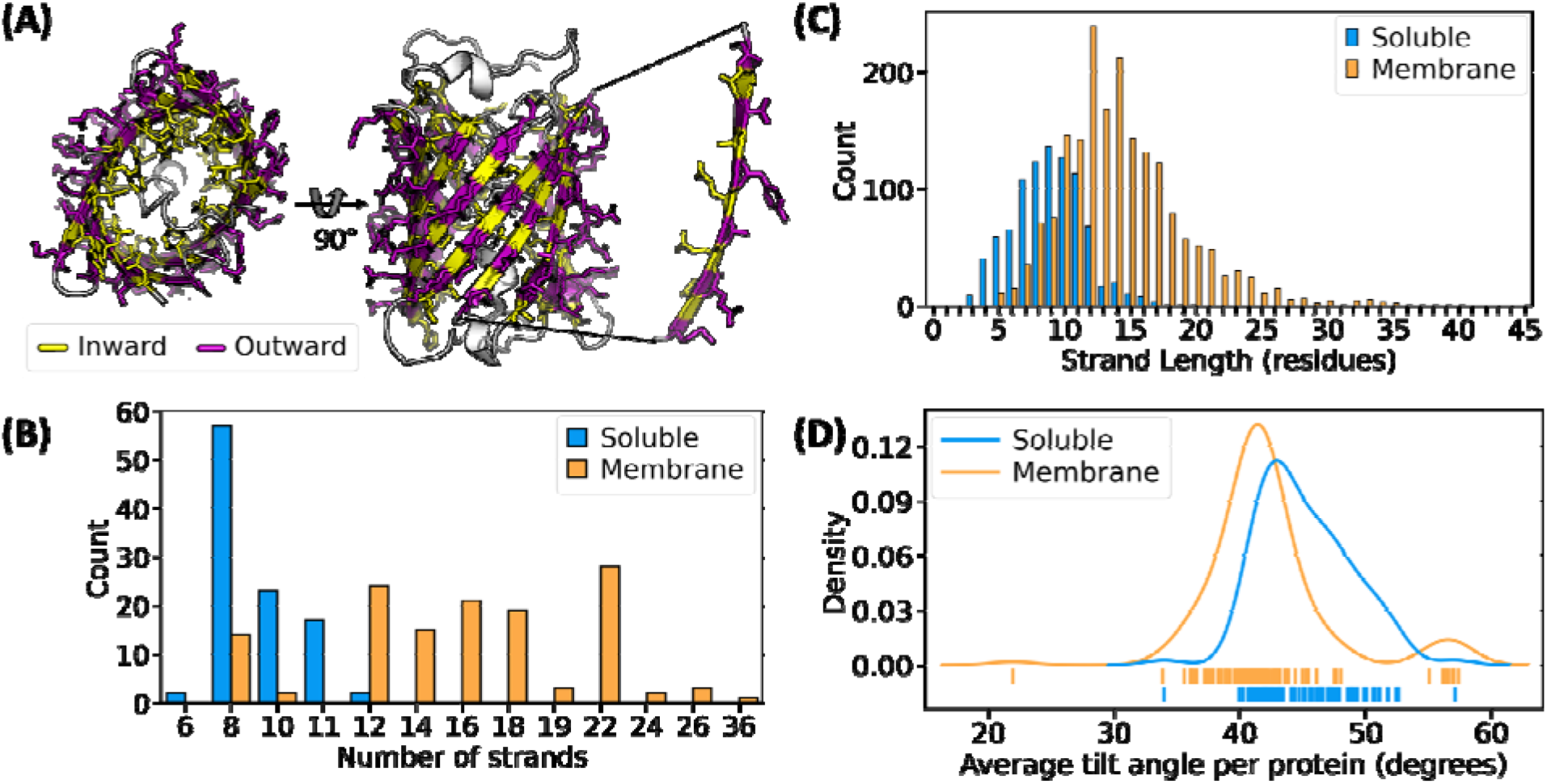
General properties of soluble and membrane barrels. (A) Structure of a soluble barrel (Green fluorescent protein FP512) with barrel residues shown as sticks, inward residues shown in yellow, outward residues shown in purple, from left to right – top view, side view, and expanded view of a single β-strand. (B-D) Soluble shown in blue, membrane shown in orange (B) number of barrels with different strand numbers, (C) histogram of number of strands with different strand lengths, (D) kernel density estimate of average tilt angle per protein, ticks at the bottom of the curve represents individual data points.

Polar Bearal was updated to identify the strands comprising the β-barrel, irrespective of sequentially contiguous strand position or strand length. For calculating the residue direction, Polar Bearal uses the angle between the axis and the vector pointing from midpoint of carboxyl carbon and nitrogen atoms of the backbone to the carbon α atom. For soluble β-barrel, this method was followed for glycine, but for all other residues it was updated to compare the distance between the nearest point on the axis and the carbon α atom to the distance of the nearest point on the axis to the carbon β atom. For membrane β-barrels, residues that were not within 13Å of the membrane center were not considered for calculations of amino acid preference, solvent accessibility, or directional/hydrophobicity alternation.

### Data analysis and graphs

Data was analyzed in Python 3.7 using the Pandas 1.2.0^18^ and graphs were generated using Seaborn v0.11.0^19^ and Matplotlib v3.3.3.^20^

### Amino acid composition

Amino acid composition was calculated for each β-barrel by looking at the frequency of each of the native 20 amino acids in either the complete β-barrel, or just the inward-facing side or outward-facing side of that β-barrel. The frequency *f*_*i*_ for each the native amino acid *i*, was calculated by counting the number of that amino acid *n*_*i*_ and dividing it with total number of residues.

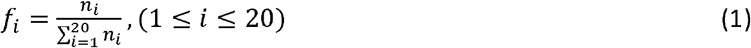

The frequency for each amino acid per β-barrel was compared between membrane β-barrels and soluble β-barrels. For computing the statistical significance, an independent two-sample t-test was performed using the SciPy library^21^ and the p-values were adjusted with Bonferroni correction using statsmodels Python module.^22^ The results were then ordered on the basis of difference between the mean amino acid frequency per protein of membrane β-barrels to that of soluble β-barrels.

To compare the hydrophobicity between soluble β-barrels and membrane β-barrels, the Kyte-Doolittle hydrophobicity scale^23^ was used. For each β-barrel, the mean hydrophobicity index *H* was calculated for the complete β-barrel, the inward-facing side and the outward-facing side, by multiplying the frequency of each amino acid *f*_*i*_ in each region to its hydrophobicity index *h*_*i*_ and then adding them up.

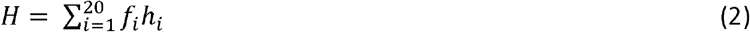

### Relative accessible surface area

Solvent accessible surface area (SASA) was calculated using FreeSASA.^24^ To calculate the SASA we used PDB files with their biological assemblies estimated through PISA.^25^ The structures are stripped off water, ligands, and metal ions by FreeSASA before SASA estimation. Relative accessible surface area for each residue in each β-barrel was calculated by dividing the SASA value by the residue’s theoretical maximum accessible surface area.^26^ Buried residues were defined as residues having <15% relative accessible surface area and exposed ≥15% relative accessible surface area.

### Alternation and alternation efficiency

Alternation of residue direction for each strand in the membrane or soluble dataset, was calculated by first recording the length of sequences within each strand where residues were facing a different direction than the previous residue. To normalize this for a subsequence (sequence fragment) of length ℓ, we divided the number of subsequences of length ℓ that were of alternating direction by the number of subsequences of length ℓ that were examined (Figure S2). For polymeric membrane β-barrels, we only examined strands from the first chain, or first two chains of PDB IDs 4TW1 and 3B07, as the other strands are redundant. We used the same calculation for alternating hydrophobicity, and consider D, E, K, N, Q, R, S, T, C, and H to be polar amino acids and A, F, I, L, V, W, Y, M, and P to be non-polar amino acids. Glycine was considered to have different hydrophobicity than any non-glycine amino acid before or after it (i.e. “DG” and “GA” are examples of alternating hydrophobicity, but “GG” is not).

## Results

In order to understand the difference between barrels in the membrane and soluble environments, we first compared the shape and size between the soluble and membrane β-barrels. We found soluble β-barrels to be smaller in size compared to membrane β-barrels, both in strand number (Figure 1B) and in strand length (Figure 1C). When controlling for strand tilt, strand length is correlated with barrel height and because strand number is correlated to the barrel circumference, strand number is linearly correlated with barrel diameter. Soluble β-barrels have strand numbers ranging from six to twelve strands. Eight-stranded soluble β-barrels are the most frequently structurally resolved soluble barrels and include lipocalins, avidins, and allene oxidase cyclases. The next most common are ten-stranded soluble lipocalin barrels followed by eleven-stranded green fluorescent protein barrels. For membrane β-barrels the number of strands in a barrel ranges from 8 strands to 36 strands in our dataset. Whereas all of the soluble barrels in our dataset are made out of single polypeptide chain, we included membrane β-barrels composed of both single and multiple polypeptide chains. The mode membrane strand number is 22, which are corked barrels like FhuA.

Membrane barrels tend to have longer strands than soluble barrels (Figure 1C). The mode soluble β-barrel strand-length is nine residues, whereas the mode membrane β-barrel strand-length is twelve residues. The few membrane strands above 30 residues are in lysins which are known to be taller.^27^ Tilt angles tend to be more similar between membrane barrels and soluble barrels (Figure 1D). Membrane barrels are slightly less tilted than soluble barrels which in addition to the strand length contributes to overall taller barrels. The exception to this is the efflux pump barrels which have tilt angles well above 50°, which is likely a result of their different evolutionary lineage^3a^.

Membrane β-barrels natively fold in a hydrophobic milieu and soluble β-barrels in a hydrophilic milieu. It is therefore understood that the exterior of soluble barrels is hydrophilic, and the exterior of membrane β-barrels are hydrophobic. Moreover, the larger strand numbers of membrane β-barrels correlate with larger barrel diameters and therefore often have solvated pores. The smaller strand numbers of soluble β-barrels translate to better-packed interiors that are not accessible to water. We calculated the mean hydrophobicity index per protein and plotted kernel density estimates of the distribution of protein hydrophobicity for soluble and membrane barrels. The overall hydrophobicity of membrane and soluble barrels are the same (Figure 2A). When the hydrophobicity is divided into the contributions of inward-facing and outward-facing residues (Figure 2B), it becomes apparent that membrane barrels are inside-out soluble barrels. The outside of membrane barrels is similarly hydrophobic to the hydrophobic core of soluble barrels and the inside of membrane barrels are slightly more hydrophilic than the outside of soluble barrel.

**Figure 2:**
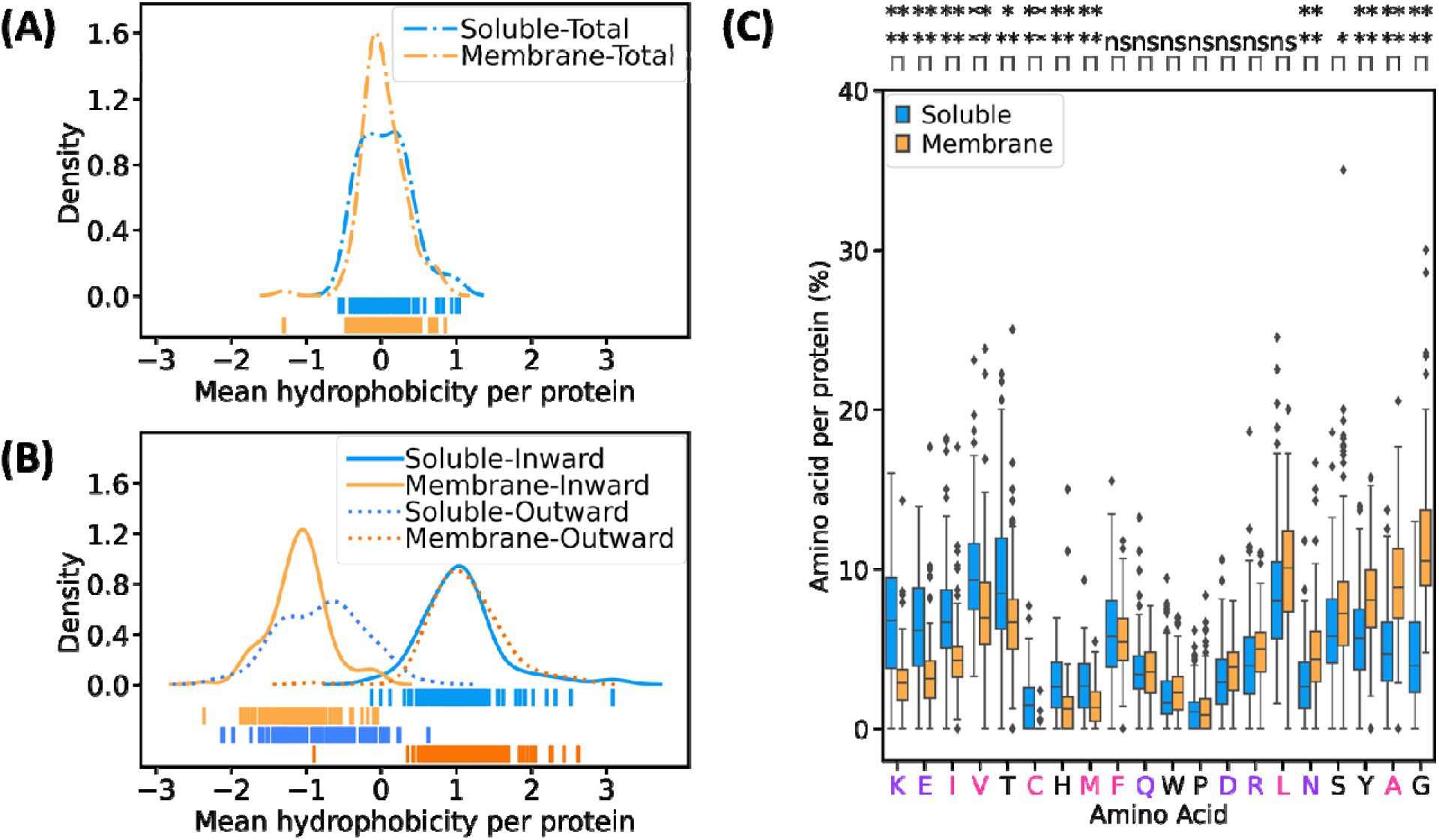
Amino acid preferences in barrels. (A) Kernel density estimate of mean hydrophobicity index per protein. Soluble shown in blue, membrane shown in orange, ticks at the bottom represents individual protein’s mean hydrophobicity index, (B) inward residues of soluble shown in lighter blue, inward residues of membrane shown in lighter orange, outward residues of soluble shown in darker blue, outward residues of membrane shown in darker orange, kernel density estimate of mean hydrophobicity index of each side (inward vs outward) per protein, ticks at the bottom represents individual protein side mean hydrophobicity index, (C) amino acid composition per protein for each dataset - soluble shown in blue, membrane shown in orange, non-polar amino acid shown in pink, polar amino acid shown in purple, box and whisker plot represents distribution of amino acid percentage in each protein, the box shows the interquartile range (IQR) with a line inside the box representing the median, the whiskers extends to 1.5 x IQR beyond upper or lower quartile, the outliers beyond the whiskers’ range are shown in black diamonds, amino acids on x-axis are arranged based on the difference between the mean amino acid percentage of soluble and the mean amino acid percentage of membrane, left is soluble preferred, right is membrane preferred, p-value is shown on top (ns: 0.05 < p ≤ 1, : 1.00e-02 < p ≤ 0.5, : 1.00e-03 < p ≤ 1.00e-02, :1.00e-04 < p ≤ 1.00e-03, : p ≤ 1.00e-04).

Different amino acids are more common in different secondary structures and the differences depend on the ϕ and ψ angles.^28^ Therefore, we compared the amino acid preferences among membrane and soluble barrels (Figure 2C, S3, and Tables ST3-7). The propensity of a few amino acids is significantly different when comparing between the complete barrels. Glycines and alanines are preferred more in membrane β-barrel than soluble β-barrel (11.6% vs 4.5% and 9.1% vs 5.9% respectively) whereas lysines and glutamates are preferred more in soluble β-barrel than membrane β-barrel (6.8% vs 2.7% and 6.3% vs 3.3% respectively). Although the soluble β-barrel core and the membrane surface are both hydrophobic, there are subtle differences as well (Figure S3C, Table ST6). In the hydrophobic regions, soluble β-barrels have a higher preference than membrane β-barrels for isoleucine, a hydrophobic β-branched amino acid long known to be favored in β-strands. This trend is not apparent for leucine— which while less hydrophobic is more lipophilic.

The phenomenon that outward-facing residues on soluble β-barrels are slightly more hydrophilic than inward-facing residues on membrane barrels (Figure 2B) may be attributed to additional helical structure outside of some β-soluble barrels forming hydrophobic contacts on the outside of soluble β-barrels. To determine the relative rates of burial of soluble and membrane β-barrels and how this contributes to mean hydrophobicity, we calculated the relative accessible surface area of each residue for membrane and soluble β-barrels (Figure 3A). For both soluble and membrane β-barrels, the outward residues are more surface accessible compared to the inward residues. However, for both inward and outward-facing residues, soluble β-barrels are less exposed than membrane β-barrels (Figure 3B). Next we calculated the mean hydrophobicity index of the buried (Figure 3C) and exposed residues (Figure 3D). Buried residues on the outward side of the barrel have similar hydrophobicities between membrane and soluble β-barrels. In contrast, buried inward residues have opposite hydrophobicities between membrane and soluble β-barrels. Soluble β-barrel, inward, buried residues are hydrophobic because they are part of a hydrophobic core. Membrane β-barrel, inward buried residues are hydrophilic not just because the middle of the barrel is often a pore, but even when it is too narrow for water accessibility the core can be held together by salt bridges.^31^ With respect to exposed residues, outward-facing, membrane β-barrel residues are hydrophobic due to their lipid environment and outward-facing soluble β-barrel residues are hydrophilic due to their aqueous environment. Exposed, inward-facing residues of membrane β-barrels are hydrophilic due to their aqueous pores. Exposed, inward-facing soluble β-barrel residues show a neutral hydrophobic tendency. While exposed residues are generally hydrophilic, the neutral trend in soluble β-barrel is likely due to residues that interact with the ligands or prosthetic groups, as many of the soluble proteins in our dataset bind hydrophobic molecules and in our analysis we removed the ligands from the structure, exposing these residues.

**Figure 3:**
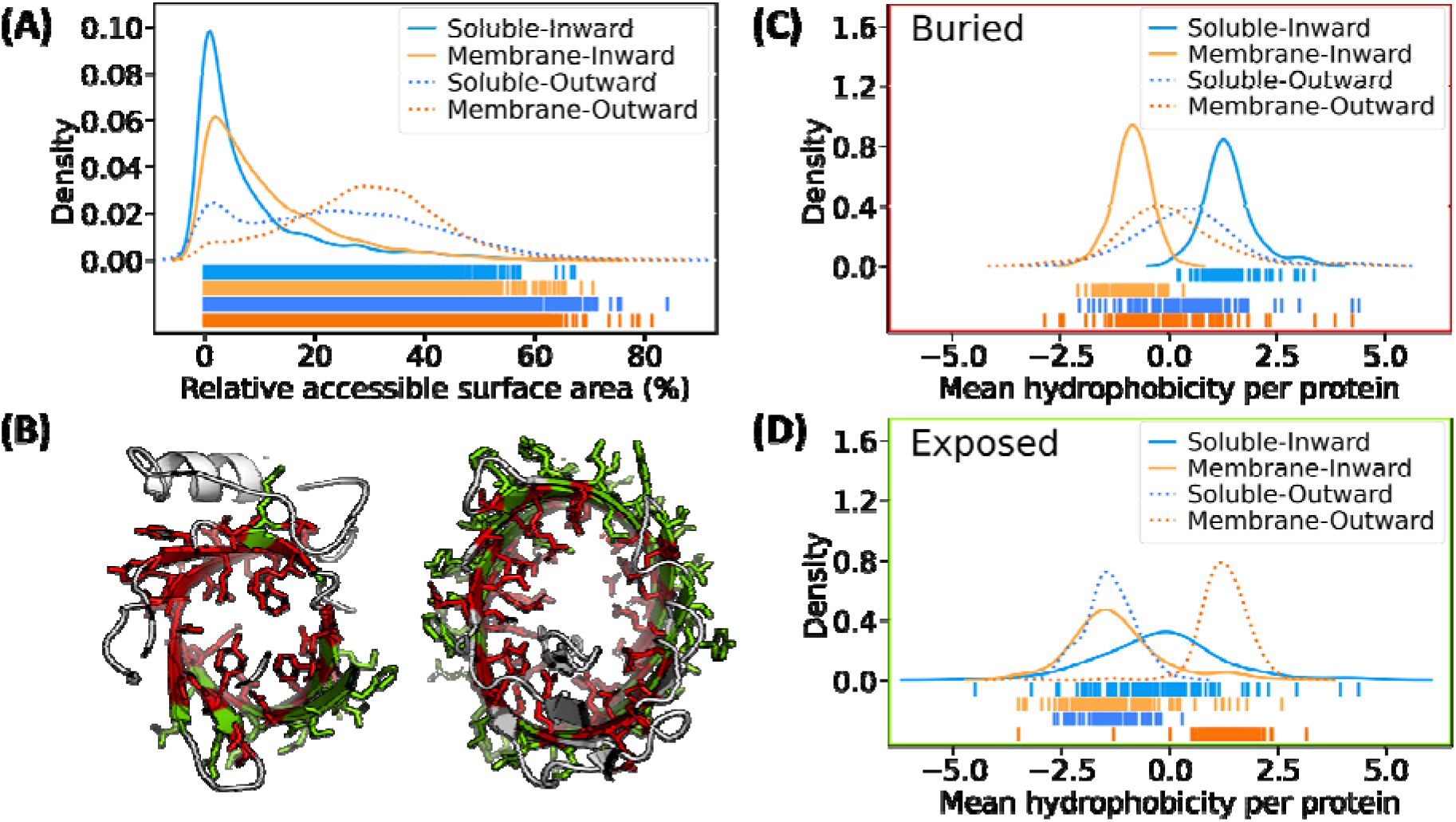
Solvent accessibility of soluble and membrane β-barrels: (A) Kernel density estimate of relative accessible surface area of each residue in membrane and soluble dataset subdivided into inward or outward direction, ticks below the curve represents individual relative accessible surface area of each residue, (A, C, D) inward residues of soluble shown in blue, inward residues of membrane shown in orange, outward residues of soluble shown in darker blue, outward residues of membrane shown in darker orange, (B) crystal structure of a soluble β-barrel female-specific histamine-binding protein 2 PDBID: 1QFT (left) and membrane β-barrel transmembrane domain of intimin PDBID: 4E1S (right) with barrel residues shown in sticks, buried residues (relative accessible surface area < 15%) are shown in red, exposed residues (relative accessible surface area ≥ 15%) are shown in green, (C-D) kernel density estimate of mean hydrophobicity per protein for (C) buried residues, (D) exposed residues, ticks below the curves represents individual mean hydrophobicity index of each protein region.

In an idealized β-strand, residues alternate consistently between facing inward and outward (Figure 1D), Tied to this understanding has been the existence of a hydrophobicity alternation in membrane β-barrels with inward residues hydrophilic and outward residues hydrophobic.^32^ To assess the degree of directional and hydrophobicity alternation in soluble and membrane β-barrels we calculated the frequency of alternating subsequences (Figure 4). Overall, the longer the subsequence the less likely the sequence is to perfectly alternate. This is to be expected. If each state were equally likely, then a two-residue subsequence would perfectly alternate PN (<u>p</u>olar <u>n</u>on-polar) or NP 50% of the time and not alternate PP or NN 50% of the time. Whereas a three-residue subsequence would only alternate 25% of the time. We find that structural alternation is very high for membrane β-barrels. Over 90% of membrane β-strands have perfect alternation when the subsequence is seven residues or fewer (Figure 4A). Soluble β-barrels have less directional alternation though soluble directional alternation is more reliable than hydrophobicity alternation (Figure 4B). We find that membrane-strand hydrophobicity alternates with ~74% efficiency and soluble strand hydrophobicity alternates at ~64% efficiency. Because the directional alternation is so much more efficient than the hydrophobicity alternation, percentage efficiencies for coupled directional and hydrophobicity alternations are similar to hydrophobicity alternations.

**Figure 4:**
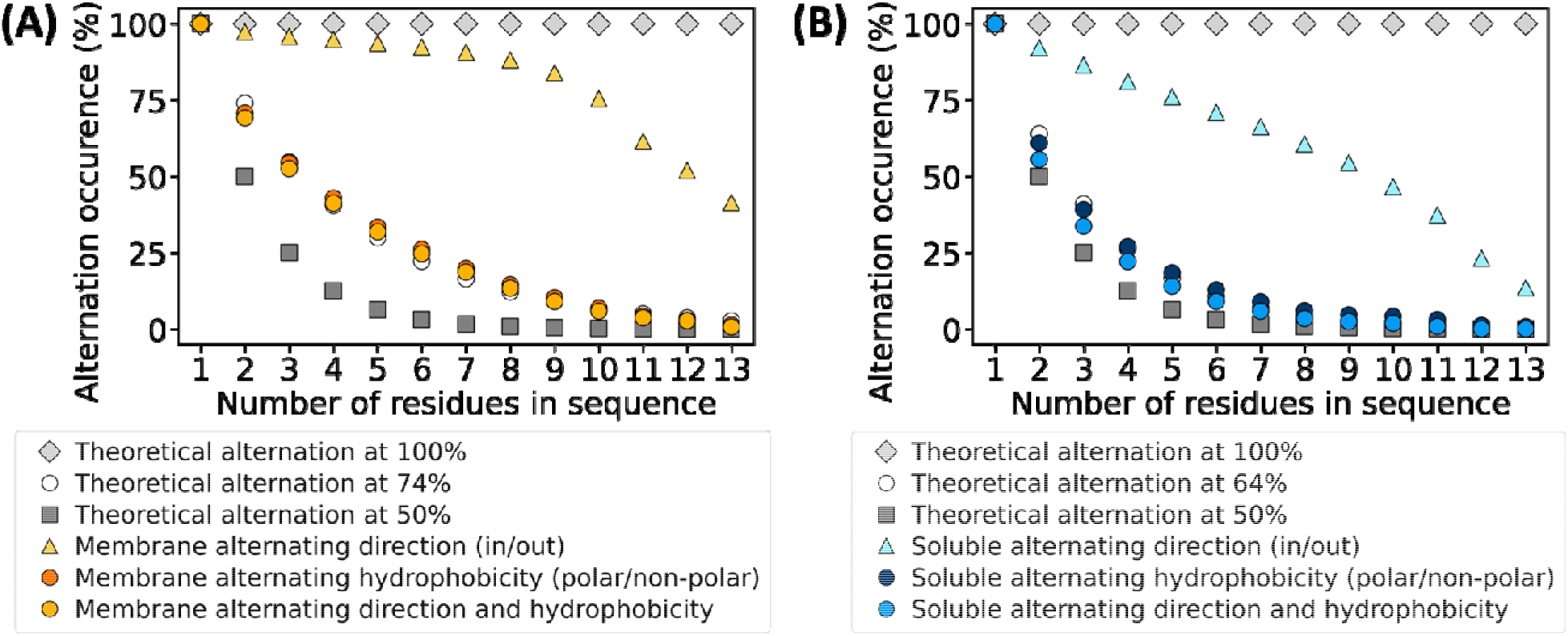
Alternation of direction and hydrophobicity of residues in soluble and membrane barrels: (A-B) Ideal alternation sequence alternating at 100% shown in grey diamond, random alternation sequence alternating at 50% shown in dark grey square, actual alternating direction shown in triangle, actual alternating hydrophobicity shown in dark colored circle, actual alternating direction and hydrophobicity shown in light colored circle, (A) membrane β-barrel shown in shades of orange, closest approximation of alternation is at efficiency of 74% shown with white circle, (B) soluble β-barrel shown in shades of blue, closest approximation of alternation is at efficiency of 64% shown with white circle.

## Discussion

Here we have compared features of membrane β-barrels to structurally similar soluble β-barrels to determine differences that facilitate the success of membrane β-barrels’ folding and functioning in the membrane milieu.

Features such as outward-facing charged residues in membrane β-barrels have been shown to be dependent on membrane height. The membrane height also creates a minimum height for membrane barrels as the loops connecting the strands must remain outside of the membrane. This constraint appears to have translated into longer strand lengths for membrane β-barrels in comparison to soluble β-barrels.

Membrane helical proteins are known to have hydrophobic cores with similar hydrophobicity to the hydrophobic cores of soluble proteins.^34^ However, membrane β-barrels have hydrophilic cores. The precision of our measurements of inside-out hydrophobicity plots between membrane and soluble β-barrels had not previously been established but the trend has long been known. Not only do soluble proteins generally have hydrophobic cores but the hydrophobic cores of many soluble barrels are used in their function of binding hydrophobic molecules and protecting sensitive fluorophores. The soluble cores of membrane β-barrels have also been known as these pores often transport polar solutes across the membrane. The use of membrane β-barrels for import and export is likely also linked to the evolution of barrels with more strands that could in turn accommodate the passage of larger molecules. In contrast, the functions of soluble β-barrels favor narrower barrels and therefore fewer strands per barrel.

Amino acid preferences like glycine are known to be used for tight packing of helical membrane proteins^35^ ^36^ and weak hydrogen bonds with the acidic carbon alpha of glycine.^37^ Though it remains unclear what its role is in membrane β-barrels, some have suggested it is required for the larger bend angles inherent in membrane β-barrels.^13^ We anticipate that the flexibility imparted by glycine and to a lesser extent alanine may also facilitate the complex process of membrane β-barrel insertion and folding into the membrane.

Though the outward regions of membrane β-barrels are as hydrophobic as the inward hydrophobic cores of soluble β-barrels, the inward hydrophilic pores of membrane β-barrels are slightly more hydrophilic than the hydrophilic surfaces of soluble β-barrels. This difference is likely due to exterior helical domains on the soluble barrels making some positions on the barrel surface buried in an intra-protein interaction. Such domains are not found on membrane barrels and are in fact likely precluded by the biological insertion mechanisms of the membrane. The inner membrane has a translocon allowing for the gating of the hydrophobic helices into the membrane^38^, and the outer membrane has the β-assembly machinery that inserts β-barrels.^39^ Though hydrophilic helices exist as plugs or periplasmic and extracellular domains to membrane barrels, it seems there is no mechanism for inserting hydrophobic helices along with membrane barrels.

Finally, although nothing in nature is perfect, soluble barrels deviate from the expected alternation pattern more than membrane barrels. One correlation of note is that membrane barrels alternate hydrophobicity with 74% efficiency and have ~74% outward-facing hydrophobic amino acids. Soluble barrels alternate hydrophobicity with 64% efficiency and have ~67% inward facing hydrophobic amino acids. Therefore, it may be that the alternation is capped by the hydrophobic content of the hydrophobic face of the barrel. With respect to the lower alternation efficiency of soluble barrels, we anticipate that perhaps the smaller size, and a greater variety of other protein domains which interact with soluble barrel regions that may contribute to these imperfections.

## Conclusions

We investigated the differences between soluble and membrane β-barrels to determine what features might be specific to the maintenance of the β-barrel fold in the membrane. We observed membrane β-barrels to be generally larger in strand number and length than soluble β-barrels. Membrane β-barrels tend to have more glycines and alanines than soluble β-barrels which tend to have more lysines and glutamates. Furthermore, the overall hydrophobicity is similar for membrane β-barrels and soluble β-barrels, but when looking at the inward and outward region separately, they have opposite hydrophobicity. Finally, we measured the periodicity of the hydrophobicity alternation in these β-barrel proteins and found membrane β-barrel alternate at better efficiency than soluble β-barrels.

## Acknowledgements

We are grateful for inspiration from Ruth Nussinov on everything from DNA sequence analysis to RNA structure to protein interactions, and we wish her a very happy birthday. This manuscript benefited from conversations with Meghan Franklin, Daniel Montezano, and Jaden Anderson as well as thoughts at the early stages of the work contributed by Pushpa Itagi and early set up of the dataset from Jaden Anderson. Funding for this work was provided by NIH awards DP2GM128201 and The Gordon and Betty Moore Inventor Fellowship.

