## Supplementary Figure and Data for "Membrane barrels are taller, fatter, inside-out soluble barrels" (from another version of this preprint: version 1, 2021-02-01)

### Supplementary Figures and Tables

#### Figure S1

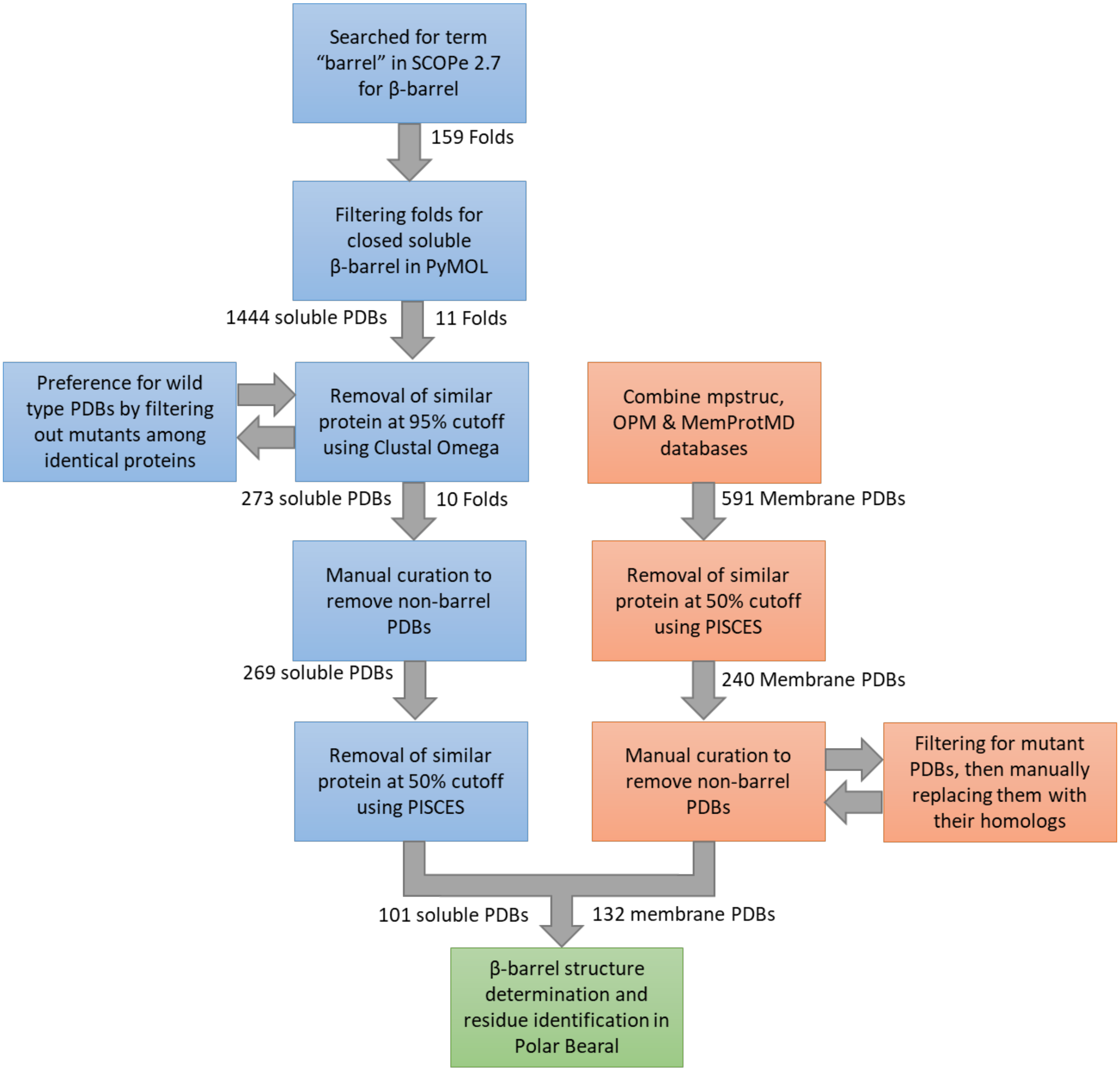


**Figure S1: Pipeline for dataset generation:** Steps for compiling the soluble β-barrel dataset shown in blue boxes, steps for compiling membrane β-barrel shown in orange boxes, common steps for both membrane and soluble β-barrel shown in green boxes, text next to grey arrow shows the number PDBs after each step.

### Figure S2

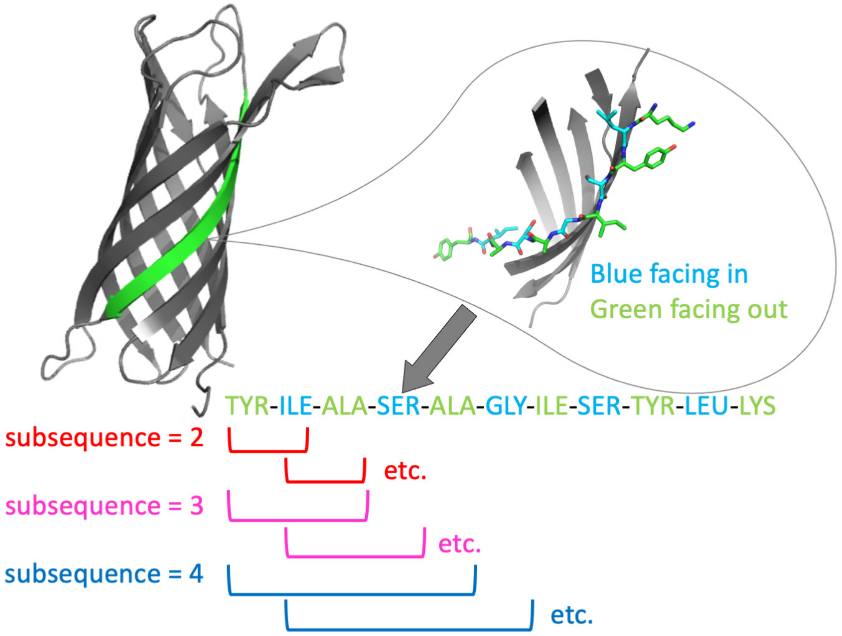


**Figure S2 Example of subsequences in alternation β-barrel:** Membrane β-barrel NspA, PDB ID - 1P4T shown in grey with green strand (left) blown up to stick view (right) with residues alternating directions shown in blue for inward direction and green for outward direction, below the sequence are examples of subsequences of length ℓ = 2, 3, and 4.

### Figure S3

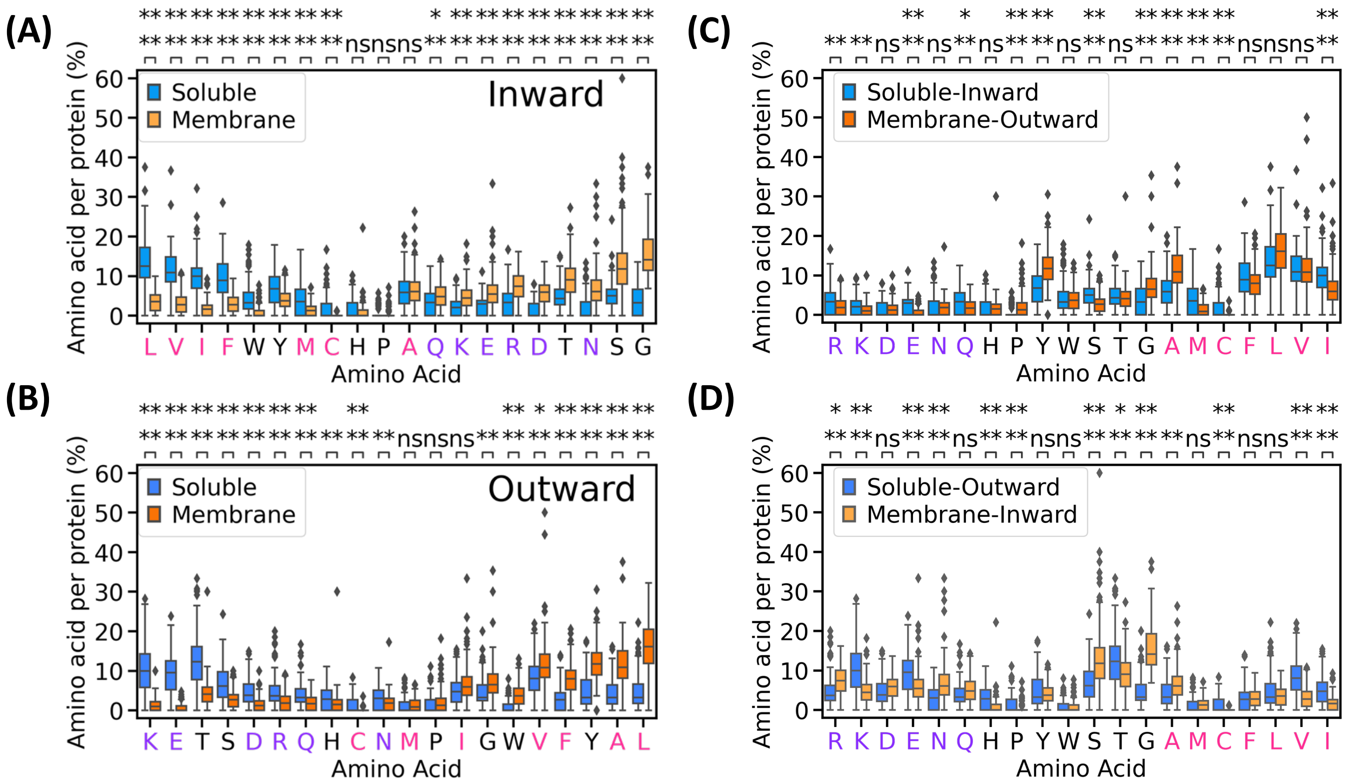


**Figure S3 Amino acid composition comparison between each β-barrel regions:** (A-D) Amino acid composition per protein for each dataset - soluble shown in blue, membrane shown in orange, non-polar amino acid shown in pink, polar amino acid shown in purple, inward residues of soluble shown in lighter blue, inward residues of membrane shown in lighter orange, outward residues of soluble shown in darker blue, outward residues of membrane shown in darker orange, box and whisker plot represents distribution of amino acid percentage in each protein, the box shows the interquartile range (IQR) with a line inside the box representing the median, the whiskers extends to 1.5 x IQR beyond upper or lower quartile, the outliers beyond the whiskers are shown in black diamond, (A-B) amino acids on x-axis are arranged based on the difference between the mean amino acid percentage of soluble and the mean amino acid percentage of membrane, (A) soluble and membrane inward-facing residues, (B) soluble and membrane outward-facing residues, (C-D) amino acids on x-axis are arranged based on their mean hydrophobic index value, left is soluble preferred, right is membrane preferred, (C) hydrophobic preferred positions, inward soluble and outward membrane, (D) hydrophilic preferred positions, outward soluble and inward membrane.

p-value is shown on top (ns: 0.05 < p ≤ 1, * : 1.00e-02 < p ≤ 0.5, ** : 1.00e-03 < p ≤ 1.00e-02, *** :1.00e-04 < p ≤ 1.00e-03, **** : p ≤ 1.00e-04).

**Table ST1**. Soluble dataset:

| PDB ID | Name | SCOPe fold | Strand Number | Biological Assembly |
|---|---|---|---|---|
| **1VKY** | S-adenosylmethionine:tRNA ribosyltransferase-isomerase (QueA) | e.53 (QueA-like) | 6 | 1 |
| **1XQB** | YaeB | e.56 (YaeB-like) | 6 | 1 |
| **2OOJ** | Hypothetical protein (NP_717203.1) | b.159 (AOC barrel-like) | 8 | 1 |
| **2Q03** | Uncharacterized protein (YP_563039.1) | b.159 (AOC barrel-like) | 8 | 1 |
| **4H6B** | Allene oxide cyclization in moss | b.159 (AOC barrel-like) | 8 | 1 |
| **2F9H** | PTS System, IIA Component | b.161 (PTSIIA/GutA-like) | 8 | 2 |
| **1AQB** | Retinol-Binding Protein | b.60 (Lipocalins) | 8 | 2 |
| **1AVG** | Triabin | b.60 (Lipocalins) | 8 | 2 |
| **1D2U** | Nitrophorin 4 | b.60 (Lipocalins) | 8 | 1 |
| **1DZJ** | Odorant-Binding Protein | b.60 (Lipocalins) | 8 | 1 |
| **1E5P** | Aphrodisin | b.60 (Lipocalins) | 8 | 1 |
| **1EPA** | Epididymal retinoic acid-binding protein | b.60 (Lipocalins) | 8 | 1 |
| **1EW3** | Allergen Equ C 1 | b.60 (Lipocalins) | 8 | 4 |
| **1LKE** | DigA16 | b.60 (Lipocalins) | 8 | 1 |
| **1QFT** | Female-Specific Histamine Binding Protein 2 | b.60 (Lipocalins) | 8 | 1 |
| **1QWD** | Lipocalin Blc | b.60 (Lipocalins) | 8 | 1 |
| **1XKI** | Von Ebner's Gland Protein | b.60 (Lipocalins) | 8 | 4 |
| **2HLV** | Odorant-Binding Protein | b.60 (Lipocalins) | 8 | 2 |
| **2HZQ** | Apolipoprotein D | b.60 (Lipocalins) | 8 | 1 |
| **2JOZ** | Hypothetical Protein YxeF | b.60 (Lipocalins) | 8 | 3 |
| **2L5P** | Lipocalin 12 | b.60 (Lipocalins) | 8 | 2 |
| **2R73** | Possum Milk Whey Lipocalin Trichosurin | b.60 (Lipocalins) | 8 | 1 |
| **2RD7** | Complement component C8 gamma chain | b.60 (Lipocalins) | 8 | 2 |
| **2WWP** | Prostaglandin-H2 D-Isomerase | b.60 (Lipocalins) | 8 | 1 |
| **2XST** | Lipocalin 15 | b.60 (Lipocalins) | 8 | 1 |
| **3FIQ** | Odorant-Binding Protein 1F | b.60 (Lipocalins) | 8 | 1 |
| **3KFF** | Major Urinary Protein 4 | b.60 (Lipocalins) | 8 | 1 |
| **3KZA** | Gyuba | b.60 (Lipocalins) | 8 | 1 |
| **3L4R** | Minor allergen Can f 2 | b.60 (Lipocalins) | 8 | 2 |
| **3S26** | Murine Siderocalin | b.60 (Lipocalins) | 8 | 2 |
| **3SAO** | Siderocalin Ex-FABP | b.60 (Lipocalins) | 8 | 1 |
| **4ALO** | H1 Crustacyanin | b.60 (Lipocalins) | 8 | 1 |

| | | | | |
|---|---|---|---|---|
| **4AW8** | Metal-Binding Protein ZinT | b.60 (Lipocalins) | 8 | 4 |
| **4ES7** | Protein AMBP | b.60 (Lipocalins) | 8 | 2 |
| **4GE1** | Biogenic Amine-Binding Protein | b.60 (Lipocalins) | 8 | 2 |
| **4ODD** | Dog lipocalin allergen | b.60 (Lipocalins) | 8 | 4 |
| **4R0B** | Glycodelin | b.60 (Lipocalins) | 8 | 1 |
| **4UU3** | Ferulic acid decarboxylase | b.60 (Lipocalins) | 8 | 2 |
| **4WFU** | Bovine allergen Bos d 2 | b.60 (Lipocalins) | 8 | 2 |
| **4XMC** | Nitrophorin-7 | b.60 (Lipocalins) | 8 | 4 |
| **5EZ2** | Sandercyanin Fluorescent Protein (SFP) | b.60 (Lipocalins) | 8 | 1 |
| **1EI5** | D-Aminopeptidase | b.61 (Streptavidin-like) | 8 | 2 |
| **1JIW** | Alkaline proteinase inhibitor APRin | b.61 (Streptavidin-like) | 8 | 2 |
| **1MK5** | Streptavidin | b.61 (Streptavidin-like) | 8 | 1 |
| **1NYC** | Cysteine Protease Inhibitor | b.61 (Streptavidin-like) | 8 | 4 |
| **1PBY** | Quinohemoprotein amine dehydrogenase 60 kda subunit | b.61 (Streptavidin-like) | 8 | 4 |
| **1Y0G** | YceI | b.61 (Streptavidin-like) | 8 | 1 |
| **2FHL** | Avidin-related protein 4/5 | b.61 (Streptavidin-like) | 8 | 1 |
| **2Y32** | Fradavidin | b.61 (Streptavidin-like) | 8 | 1 |
| **2ZSC** | Tamavidin2 | b.61 (Streptavidin-like) | 8 | 1 |
| **3HPE** | Acidic stress response factor HP1286 is a YceI homolog | b.61 (Streptavidin-like) | 8 | 2 |
| **3PDF** | Cathepsin C | b.61 (Streptavidin-like) | 8 | 2 |
| **3SZH** | Shwanavidin | b.61 (Streptavidin-like) | 8 | 1 |
| **4BJ8** | Zebavidin | b.61 (Streptavidin-like) | 8 | 1 |
| **4GGR** | bradavidin2 (Form A) | b.61 (Streptavidin-like) | 8 | 1 |
| **4Z6J** | Hoefavidin | b.61 (Streptavidin-like) | 8 | 2 |
| **5IXG** | BcnB | b.61 (Streptavidin-like) | 8 | 2 |
| **1JSG** | Oncogene Product P14TCL1 | b.63 (Oncogene products) | 8 | 1 |
| **1QTT** | Product of The MTCP1 Oncogene | b.63 (Oncogene products) | 8 | 1 |
| **1GGL** | Protein (Cellular Retinol-Binding Protein III) | b.60 (Lipocalins) | 10 | 4 |
| **1IFC** | Intestinal Fatty Acid Binding Protein | b.60 (Lipocalins) | 10 | 2 |
| **1LFO** | Liver Fatty Acid Binding Protein | b.60 (Lipocalins) | 10 | 4 |
| **1MDC** | Insect Fatty Acid Binding Protein | b.60 (Lipocalins) | 10 | 2 |
| **1O8V** | Fatty Acid Binding Protein Homolog | b.60 (Lipocalins) | 10 | 1 |
| **1R0U** | YwiB | b.60 (Lipocalins) | 10 | 2 |
| **1VPR** | Luciferase | b.60 (Lipocalins) | 10 | 2 |

| | | | | |
|---|---|---|---|---|
| **1VYF** | 14 Kda Fatty Acid Binding Protein | b.60 (Lipocalins) | 10 | 1 |
| **2A0A** | Der F 13 | b.60 (Lipocalins) | 10 | 1 |
| **2FR2** | Hypothetical Protein Rv2717C | b.60 (Lipocalins) | 10 | 1 |
| **2N93** | lcFABP | b.60 (Lipocalins) | 10 | 1 |
| **2O62** | Hypothetical protein (ZP_00105914.2) | b.60 (Lipocalins) | 10 | 4 |
| **2QO4** | Liver-Basic Fatty Acid Binding Protein | b.60 (Lipocalins) | 10 | 2 |
| **3EMM** | AT1G79260 | b.60 (Lipocalins) | 10 | 1 |
| **3IA8** | THAP Domain-Containing Protein 4 | b.60 (Lipocalins) | 10 | 1 |
| **3P6D** | atty-acid binding protein (FABP4) | b.60 (Lipocalins) | 10 | 1 |
| **3PPT** | REP1-NXSQ fatty acid transporter | b.60 (Lipocalins) | 10 | 2 |
| **4AZP** | epidermal fatty acid-binding protein (FABP5) | b.60 (Lipocalins) | 10 | 2 |
| **4I3B** | Bilirubin-inducible fluorescent protein UnaG | b.60 (Lipocalins) | 10 | 2 |
| **5GGE** | Fatty Acid Bindin Protein, Isoform B | b.60 (Lipocalins) | 10 | 4 |
| **5HZQ** | Cellular Retinoic Acid-Binding Protein 2 | b.60 (Lipocalins) | 10 | 4 |
| **5L8I** | Fatty Acid Binding Protein 6 | b.60 (Lipocalins) | 10 | 4 |
| **6AT8** | Retinoid-Binding Protein 7 | b.60 (Lipocalins) | 10 | 1 |
| **2ICH** | Putative AttH | b.176 (AttH-like) | 11 | 1 |
| **1GL4** | Nidogen-1 | d.22 (GFP-like) | 11 | 2 |
| **2C9J** | Fluorescent protein CMFP512 | d.22 (GFP-like) | 11 | 3 |
| **2DD7** | GFP-like protein | d.22 (GFP-like) | 11 | 2 |
| **2G3O** | copGFP | d.22 (GFP-like) | 11 | 4 |
| **2GW3** | Kaede | d.22 (GFP-like) | 11 | 1 |
| **2H5O** | mOrange | d.22 (GFP-like) | 11 | 1 |
| **2HPW** | Green Fluorescent Protein | d.22 (GFP-like) | 11 | 2 |
| **2ICR** | zRFP574 | d.22 (GFP-like) | 11 | 1 |
| **2RH7** | GFP | d.22 (GFP-like) | 11 | 2 |
| **2WIQ** | Killerred | d.22 (GFP-like) | 11 | 1 |
| **2ZO6** | Cyan-Emitting GFP-Like Protein, Kusabira-Cyan (KCY) | d.22 (GFP-like) | 11 | 1 |
| **3PIB** | eqFP578 | d.22 (GFP-like) | 11 | 2 |
| **4DKN** | GFPA1 | d.22 (GFP-like) | 11 | 4 |
| **4HE4** | phiYFP | d.22 (GFP-like) | 11 | 1 |
| **4HVF** | lanGFP | d.22 (GFP-like) | 11 | 1 |
| **4RYS** | NowGFP | d.22 (GFP-like) | 11 | 2 |

| | | | | |
|---|---|---|---|---|
| **1ZXU** | AT5G01750 | d.23 (Tubby C-terminal domain-like) | 12 | 1 |
| **2FIM** | Tubby related protein 1 | d.23 (Tubby C-terminal domain-like) | 12 | 1 |

**Table ST2**. Membrane Dataset:

| PDB ID | Name | SCOPe fold | Strand Number | Biological Assembly | Mono / Poly |
|---|---|---|---|---|---|
| **1P4T** | Outer Membrane Protein NspA | f.4 (Transmembrane beta-barrels) | 8 | 1 | mono |
| **1QJ8** | Outer Membrane Protein X | f.4 (Transmembrane beta-barrels) | 8 | 1 | mono |
| **1QJP** | Outer Membrane Protein A | f.4 (Transmembrane beta-barrels) | 8 | 1 | mono |
| **2ERV** | Outer membrane enzyme PagL | Not Classified | 8 | 1 | mono |
| **2F1V** | Outer Membrane Protein W | Not Classified | 8 | 3 | mono |
| **2LHF** | Outer Membrane Protein H1 | Not Classified | 8 | 1 | mono |
| **2MAF** | Opacity Protein Opa60 | Not Classified | 8 | 1 | mono |
| **2X27** | Outer Membrane Protein OprG | Not Classified | 8 | 1 | mono |
| **3DZM** | Hypothetical Conserved Protein HB27 | Not Classified | 8 | 1 | mono |
| **3GP6** | Protein PagP | f.4 (Transmembrane beta-barrels) | 8 | 1 | mono |
| **3QRA** | Attachment Invasion Locus Protein | f.4 (Transmembrane beta-barrels) | 8 | 1 | mono |
| **4FUV** | CarO | Not Classified | 8 | 1 | mono |
| **4RLC** | Outer Membrane Porin F | Not Classified | 8 | 1 | mono |
| **6QWR** | Outer Membrane Protein AlkL | Not Classified | 8 | 1 | mono |
| **2VDF** | Outer Membrane Protein OpcA | f.4 (Transmembrane beta-barrels) | 10 | 1 | mono |
| **2X55** | Plasminogen Activator Pla | f.4 (Transmembrane beta-barrels) | 10 | 1 | mono |
| **1QD5** | Outer Membrane Phospholipase A | f.4 (Transmembrane beta-barrels) | 12 | 1 | mono |
| **1TLY** | Nucleoside-Specific Channel-Forming Protein Tsx | f.4 (Transmembrane beta-barrels) | 12 | 1 | mono |
| **1UYN** | NalP | f.4 (Transmembrane beta-barrels) | 12 | 1 | mono |
| **2WJR** | Probable N-Acetylneuraminic Acid Outer Membrane Channel | Not Classified | 12 | 1 | mono |
| **3AEH** | Hemoglobin-Binding Protease Hbp Autotransporter | Not Classified | 12 | 2 | mono |

| | | | | | |
|---|---|---|---|---|---|
| **3FID** | Putative Outer Membrane Protein (LpxR) | Not Classified | 12 | 1 | mono |
| **3KVN** | Esterase EstA | Not Classified | 12 | 1 | mono |
| **3QQ2** | Brka Autotransporter | Not Classified | 12 | 1 | mono |
| **4E1S** | Intimin | Not Classified | 12 | 1 | mono |
| **4E1T** | Invasin | Not Classified | 12 | 1 | mono |
| **4FQE** | Oligogalacturonate-Specific Porin KdgM | Not Classified | 12 | 1 | mono |
| **4MEE** | Autotransporter AIDA-I | Not Classified | 12 | 1 | mono |
| **4RL8** | COG4313 outer membrane channel | Not Classified | 12 | 1 | mono |
| **5O65** | FapF | Not Classified | 12 | 3 | mono |
| **1T16** | Long-Chain Fatty Acid Transport Protein FadL | f.4 (Transmembrane beta-barrels) | 14 | 1 | mono |
| **2X9K** | Outer Membrane Protein G | Not Classified | 14 | 1 | mono |
| **3BRY** | Toluene transporter TbuX | Not Classified | 14 | 1 | mono |
| **3BS0** | Toluene transporter TodX | Not Classified | 14 | 1 | mono |
| **3DWO** | FadL homologue | Not Classified | 14 | 1 | mono |
| **4D5B** | CymA | Not Classified | 14 | 1 | mono |
| **6GIE** | Omp33 | Not Classified | 14 | 1 | mono |
| **6H3I_F** | PorV | Not Classified | 14 | 1 | mono |
| **1PRN** | Porin | f.4 (Transmembrane beta-barrels) | 16 | 3 | mono |
| **2FGR** | Outer membrane porin protein 32 | f.4 (Transmembrane beta-barrels) | 16 | 3 | mono |
| **2POR** | Porin | f.4 (Transmembrane beta-barrels) | 16 | 3 | mono |
| **3A2S** | PorB | Not Classified | 16 | 3 | mono |
| **4C00** | Translocation And Assembly Module TamA | Not Classified | 16 | 1 | mono |
| **4GEY** | Porin B | Not Classified | 16 | 1 | mono |
| **4K3B** | Outer Membrane Protein Assembly Factor BamA | Not Classified | 16 | 1 | mono |
| **4K3C** | Outer Membrane Protein Assembly Factor BamA | Not Classified | 16 | 1 | mono |
| **4N75** | Outer Membrane Protein Assembly Factor BamA | Not Classified | 16 | 1 | mono |
| **4QKY** | Filamentous Hemagglutinin Transporter Protein FhaC | Not Classified | 16 | 1 | mono |
| **4RJW** | Porin O | Not Classified | 16 | 3 | mono |
| **4Y25** | PgaA exopolysaccharide transporter | Not Classified | 16 | 1 | mono |
| **5FVN** | OmpC Porin | f.4 (Transmembrane beta-barrels) | 16 | 1 | mono |

| | | | | | |
|---|---|---|---|---|---|
| **5MDO** | Chitoporin | Not Classified | 16 | 3 | mono |
| **6EHB** | Outer Membrane Protein U | Not Classified | 16 | 3 | mono |
| **6EHD** | OmpT Protein | Not Classified | 16 | 1 | mono |
| **6EUS** | DcaP outer membrane channel | Not Classified | 16 | 3 | mono |
| **6TZK** | Cellulose Synthase Operon Protein C | Not Classified | 16 | 1 | mono |
| **1A0S** | ScrY sucrose-specific porin | f.4 (Transmembrane beta-barrels) | 18 | 3 | mono |
| **1AF6** | Maltoporin | f.4 (Transmembrane beta-barrels) | 18 | 3 | mono |
| **2YNK** | WZI | Not Classified | 18 | 1 | mono |
| **3SY7** | OprD | f.4 (Transmembrane beta-barrels) | 18 | 1 | mono |
| **3SY9** | Histidine Porin OpdC | f.4 (Transmembrane beta-barrels) | 18 | 1 | mono |
| **3SYB** | Glycine-Glutamate Dipeptide Porin OpdP | f.4 (Transmembrane beta-barrels) | 18 | 1 | mono |
| **3SYS** | Vanillate Porin OpdK | Not Classified | 18 | 1 | mono |
| **3SZV** | Pyroglutatmate Porin OpdO | f.4 (Transmembrane beta-barrels) | 18 | 1 | mono |
| **3T0S** | OccK4 (OpdL) | f.4 (Transmembrane beta-barrels) | 18 | 1 | mono |
| **4AFK** | Alginate Production Protein AlgE | Not Classified | 18 | 1 | mono |
| **4FRX** | Anaerobically-Induced Outer Membrane Porin OprE | Not Classified | 18 | 1 | mono |
| **4FSO** | OccK10 (OpdN) | f.4 (Transmembrane beta-barrels) | 18 | 1 | mono |
| **4FSP** | OccK11 (OpdR) | f.4 (Transmembrane beta-barrels) | 18 | 1 | mono |
| **5DL5** | OccAB1 | f.4 (Transmembrane beta-barrels) | 18 | 1 | mono |
| **5DL6** | Putative Porin For Vanillate Trafficking (VanP) | f.4 (Transmembrane beta-barrels) | 18 | 1 | mono |
| **5DL7** | OccAB3 | f.4 (Transmembrane beta-barrels) | 18 | 1 | mono |
| **5DL8** | Benzoate Transport Porin BenP | f.4 (Transmembrane beta-barrels) | 18 | 1 | mono |
| **5LDV** | MOMP Porin | Not Classified | 18 | 3 | mono |
| **3EMN** | Voltage-Dependent Anion-Selective Channel Protein 1 | Not Classified | 19 | 2 | mono |
| **5O8O** | Mitochondrial Import Receptor Subunit Tom40 | Not Classified | 19 | 1 | mono |

| | | | | | |
|---|---|---|---|---|---|
| **6UCU** | Mitochondrial Import Receptor Subunit Tom40 | Not Classified | 19 | 1 | mono |
| **1KMO** | Iron(III) Dicitrate Transport Protein FecA | f.4 (Transmembrane beta-barrels) | 22 | 1 | mono |
| **1XKW** | Fe(III)-Pyochelin Receptor FptA | f.4 (Transmembrane beta-barrels) | 22 | 1 | mono |
| **2GSK** | Vitamin B12 Transporter BtuB | f.4 (Transmembrane beta-barrels) | 22 | 2 | mono |
| **2HDI** | Colicin I Receptor | Not Classified | 22 | 1 | mono |
| **2IAH** | Ferripyoverdine Receptor | Not Classified | 22 | 2 | mono |
| **3CSL** | Hasr Protein,Hemophore Hasa | d.35 (Heme-binding protein A (HasA)) | 22 | 1 | mono |
| **3EFM** | Ferric Alcaligin Siderophore Receptor FauA | f.4 (Transmembrane beta-barrels) | 22 | 1 | mono |
| **3FHH** | Outer Membrane Heme Receptor ShuA | Not Classified | 22 | 1 | mono |
| **3QLB** | Enantio-Pyochelin Receptor | Not Classified | 22 | 1 | mono |
| **3V8X** | transferrin binding protein 1 TbpA | Not Classified | 22 | 1 | mono |
| **4AIP** | Fe-Regulated Protein B FrpB | Not Classified | 22 | 3 | mono |
| **4CU4** | Ferrichrome-Iron Receptor FhuA | f.4 (Transmembrane beta-barrels) | 22 | 2 | mono |
| **4EPA** | Ferric yersiniabactin uptake receptor FyuA | Not Classified | 22 | 1 | mono |
| **4RDR** | ZnuD | Not Classified | 22 | 1 | mono |
| **4ZGV** | Ferredoxin Receptor FusA | Not Classified | 22 | 1 | mono |
| **5FOK** | Iron Transport Outer Membrane Receptor PiuA | f.4 (Transmembrane beta-barrels) | 22 | 1 | mono |
| **5FP1** | Tonb-Dependent Siderophore Receptor PiuA | f.4 (Transmembrane beta-barrels) | 22 | 1 | mono |
| **5FQ8_B** | Outer Membrane Protein Omp121 | Not Classified | 22 | 1 | mono |
| **5FR8** | Tonb-Dependent Siderophore Receptor PirA | f.4 (Transmembrane beta-barrels) | 22 | 1 | mono |
| **5M9B** | Ferric Enterobactin Receptor PfeA | Not Classified | 22 | 1 | mono |
| **5T3R_D** | SusC homolog | Not Classified | 22 | 1 | mono |
| **6BPM** | Catecholate Siderophore Receptor Fiu | Not Classified | 22 | 1 | mono |
| **6E4V** | FhuE Receptor | Not Classified | 22 | 1 | mono |
| **6FOK** | Putative Copper Transport Outer Membrane Porin OprC | Not Classified | 22 | 1 | mono |

| | | | | | |
|---|---|---|---|---|---|
| **6H7F** | BauA | Not Classified | 22 | 1 | mono |
| **6I96** | Ferric Hydroxamate Uptake FoxA | Not Classified | 22 | 1 | mono |
| **6SLJ** | RagAB | Not Classified | 22 | 1 | mono |
| **6V81** | Probable TonB-Dependent Receptor YncD | Not Classified | 22 | 1 | mono |
| **3FIP** | Outer Membrane Usher Protein PapC | Not Classified | 24 | 1 | mono |
| **3OHN** | Outer Membrane Usher Protein FimD | Not Classified | 24 | 1 | mono |
| **4Q35** | Lps-Assembly Protein LptD | Not Classified | 26 | 2 | mono |
| **5IVA** | Lps-Assembly Protein LptD | Not Classified | 26 | 2 | mono |
| **6R2Q_B** | MtrAB electron transporter complex | Not Classified | 26 | 3 | mono |
| **1EK9** | Outer Membrane Protein TolC | f.5 (Outer membrane efflux proteins (OEP)) | 12 | 3 | poly |
| **1YC9** | VceC | f.5 (Outer membrane efflux proteins (OEP)) | 12 | 3 | poly |
| **2GR7** | Adhesin | Not Classified | 12 | 3 | poly |
| **2LME** | Adhesin YadA | Not Classified | 12 | 3 | poly |
| **3D5K** | Outer Membrane Protein OprM | f.5 (Outer membrane efflux proteins (OEP)) | 12 | 3 | poly |
| **4K7R** | CusC | f.5 (Outer membrane efflux proteins (OEP)) | 12 | 3 | poly |
| **4MT0** | MtrE | Not Classified | 12 | 3 | poly |
| **4MT4** | CmeC | Not Classified | 12 | 3 | poly |
| **5AZP** | Multidrug Efflux Outer Membrane Protein OprN | f.5 (Outer membrane efflux proteins (OEP)) | 12 | 3 | poly |
| **5AZS** | Outer Membrane Protein OprJ | f.5 (Outer membrane efflux proteins (OEP)) | 12 | 3 | poly |
| **3J9C** | Protective antigen PA-63 | Not Classified | 14 | 7 | poly |
| **3O44** | Hemolysin | Not Classified | 14 | 7 | poly |
| **3W9T** | Hemolytic Lectin CEL-III | d.281 (Hemolytic lectin CEL-III, C-terminal domain) | 14 | 7 | poly |
| **4H56** | Necrotic Enteritis Toxin B | Not Classified | 14 | 7 | poly |
| **6RB9** | Epsilon-Toxin Type B | Not Classified | 14 | 7 | poly |
| **6UWR** | ADP-ribosyltransferase binding component | Not Classified | 14 | 7 | poly |
| **7AHL** | Alpha-Hemolysin | f.6 (Leukocidin-like) | 14 | 7 | poly |
| **1UUN** | MspA | f.6 (Leukocidin-like) | 16 | 8 | poly |
| **3B07** | Gamma-Hemolysin Component A and B | f.6 (Leukocidin-like) | 16 | 8 | poly |
| **4TW1** | Bi-component Toxin LukGH | f.6 (Leukocidin-like) | 16 | 8 | poly |
| **5GAQ** | Lysenin | Not Classified | 18 | 9 | poly |
| **3X2R** | CsgG | Not Classified | 36 | 9 | poly |

**Table ST3:** Amino acid composition comparison per protein: Membrane β-barrel vs. soluble β-barrel. Values are shown graphically in Figure 2C.

| Amino Acid | Membrane total barrel | Soluble total barrel | Difference | Adjusted p-value |
|---|---|---|---|---|
| **ALA** | 9.10 ± 3.38 | 5.19 ± 2.86 | 3.90 | 1.97E-16 |
| **ARG** | 4.70 ± 2.37 | 4.35 ± 3.23 | 0.35 | 1.00E+00 |
| **ASN** | 4.59 ± 2.87 | 2.82 ± 2.28 | 1.76 | 1.63E-05 |
| **ASP** | 3.51 ± 1.76 | 3.19 ± 2.29 | 0.31 | 1.00E+00 |
| **CYS** | 0.04 ± 0.25 | 1.63 ± 1.60 | -1.59 | 1.95E-22 |
| **GLU** | 3.32 ± 2.39 | 6.31 ± 3.07 | -2.99 | 1.18E-13 |
| **GLN** | 3.52 ± 1.98 | 3.75 ± 2.30 | -0.24 | 1.00E+00 |
| **GLY** | 11.59 ± 4.23 | 4.46 ± 2.97 | 7.13 | 9.47E-33 |
| **HIS** | 1.41 ± 1.79 | 2.66 ± 2.02 | -1.25 | 2.36E-05 |
| **ILE** | 4.35 ± 2.40 | 7.20 ± 3.36 | -2.85 | 2.14E-11 |
| **LEU** | 9.85 ± 3.95 | 8.72 ± 4.48 | 1.12 | 8.74E-01 |
| **LYS** | 2.90 ± 2.00 | 6.84 ± 3.62 | -3.93 | 3.16E-20 |
| **MET** | 1.46 ± 1.26 | 2.60 ± 1.92 | -1.14 | 2.82E-06 |
| **PHE** | 5.55 ± 2.35 | 5.91 ± 2.95 | -0.36 | 1.00E+00 |
| **PRO** | 1.36 ± 1.64 | 1.15 ± 1.33 | 0.21 | 1.00E+00 |
| **SER** | 8.00 ± 4.60 | 6.16 ± 3.31 | 1.84 | 1.59E-02 |
| **THR** | 6.84 ± 3.28 | 9.08 ± 4.51 | -2.24 | 3.47E-04 |
| **TRP** | 2.37 ± 1.77 | 2.31 ± 2.14 | 0.05 | 1.00E+00 |
| **TYR** | 8.02 ± 3.03 | 5.77 ± 3.00 | 2.26 | 9.23E-07 |
| **VAL** | 7.53 ± 3.53 | 9.88 ± 3.57 | -2.35 | 2.18E-05 |

**Table ST4:** Amino acid composition comparison per protein region: Membrane β-barrel inward-facing vs soluble β-barrel inward-facing. Values are shown graphically in S3A.

| Amino Acid | Membrane inward | Soluble inward | Difference | Adjusted p-value |
|---|---|---|---|---|
| **ALA** | 6.42 ± 4.28 | 6.18 ± 4.29 | 0.24 | 1.00E+00 |
| **ARG** | 7.22 ± 3.91 | 3.62 ± 3.52 | 3.59 | 1.18E-10 |
| **ASN** | 7.12 ± 5.65 | 2.30 ± 3.21 | 4.82 | 9.05E-12 |
| **ASP** | 5.52 ± 3.00 | 1.42 ± 1.96 | 4.10 | 1.25E-24 |
| **CYS** | 0.03 ± 0.17 | 1.84 ± 2.94 | -1.81 | 3.77E-10 |
| **GLU** | 5.99 ± 4.64 | 2.82 ± 2.57 | 3.17 | 5.50E-08 |
| **GLN** | 5.18 ± 3.35 | 3.29 ± 2.96 | 1.89 | 2.41E-04 |
| **GLY** | 16.03 ± 6.26 | 3.90 ± 3.87 | 12.13 | 1.03E-41 |
| **HIS** | 1.01 ± 2.21 | 1.75 ± 2.36 | -0.75 | 2.81E-01 |
| **ILE** | 1.97 ± 1.93 | 9.85 ± 5.25 | -7.87 | 1.33E-37 |

| | | | | |
|---|---|---|---|---|
| **LEU** | 3.58 ± 2.74 | 13.66 ± 6.96 | -10.08 | 2.66E-35 |
| **LYS** | 4.50 ± 3.26 | 2.30 ± 2.58 | 2.21 | 1.27E-06 |
| **MET** | 1.52 ± 1.50 | 4.20 ± 3.63 | -2.68 | 9.48E-12 |
| **PHE** | 2.97 ± 2.25 | 9.31 ± 4.97 | -6.34 | 4.33E-28 |
| **PRO** | 0.46 ± 1.12 | 0.63 ± 1.31 | -0.17 | 1.00E+00 |
| **SER** | 13.25 ± 8.53 | 5.17 ± 3.79 | 8.08 | 4.12E-15 |
| **THR** | 9.35 ± 4.73 | 4.93 ± 3.41 | 4.42 | 1.77E-12 |
| **TRP** | 0.87 ± 1.39 | 4.15 ± 3.96 | -3.28 | 4.85E-15 |
| **TYR** | 3.92 ± 2.62 | 7.00 ± 4.06 | -3.08 | 5.06E-10 |
| **VAL** | 3.11 ± 2.57 | 11.69 ± 5.37 | -8.58 | 2.27E-38 |

**Table ST5:** Amino acid composition comparison per protein region: Membrane β-barrel outward-facing vs soluble β-barrel outward-facing. Values are shown graphically in S3B.

| **Amino Acid** | **Membrane outward** | **Soluble outward** | **Difference** | **Adjusted p-value** |
|---|---|---|---|---|
| **ALA** | 11.77 ± 5.67 | 4.33 ± 3.73 | 7.44 | 5.79E-23 |
| **ARG** | 2.19 ± 2.24 | 4.91 ± 4.54 | -2.72 | 1.46E-07 |
| **ASN** | 2.07 ± 2.17 | 3.28 ± 2.74 | -1.21 | 4.39E-03 |
| **ASP** | 1.52 ± 1.63 | 4.69 ± 3.92 | -3.18 | 8.02E-14 |
| **CYS** | 0.06 ± 0.36 | 1.45 ± 2.02 | -1.39 | 5.18E-12 |
| **GLU** | 0.67 ± 1.01 | 9.33 ± 5.20 | -8.66 | 8.32E-47 |
| **GLN** | 1.88 ± 1.82 | 4.12 ± 3.38 | -2.24 | 1.12E-08 |
| **GLY** | 7.21 ± 4.88 | 4.87 ± 3.90 | 2.33 | 2.19E-03 |
| **HIS** | 1.81 ± 2.94 | 3.41 ± 3.02 | -1.60 | 1.27E-03 |
| **ILE** | 6.70 ± 4.84 | 4.96 ± 3.88 | 1.74 | 6.76E-02 |
| **LEU** | 16.05 ± 6.71 | 4.59 ± 4.39 | 11.45 | 2.85E-34 |
| **LYS** | 1.31 ± 1.55 | 10.78 ± 6.53 | -9.47 | 2.96E-38 |
| **MET** | 1.40 ± 1.77 | 1.20 ± 1.85 | 0.20 | 1.00E+00 |
| **PHE** | 8.10 ± 4.15 | 3.03 ± 2.96 | 5.07 | 8.00E-20 |
| **PRO** | 2.25 ± 3.00 | 1.57 ± 2.30 | 0.68 | 1.00E+00 |
| **SER** | 2.82 ± 2.36 | 6.96 ± 4.63 | -4.14 | 3.59E-15 |
| **THR** | 4.33 ± 3.59 | 12.65 ± 7.22 | -8.32 | 2.99E-23 |
| **TRP** | 3.84 ± 2.89 | 0.80 ± 1.54 | 3.05 | 3.03E-17 |
| **TYR** | 12.06 ± 4.98 | 4.69 ± 4.25 | 7.37 | 1.47E-24 |
| **VAL** | 11.95 ± 6.84 | 8.37 ± 4.48 | 3.59 | 1.57E-04 |

**Table ST6:** Amino acid composition comparison per protein region: Membrane β-barrel inward-facing vs soluble β-barrel outward-facing. Values are shown graphically in S3C.

| **Amino Acid** | **Membrane inward** | **Soluble outward** | **Difference** | **Adjusted p-value** |
|---|---|---|---|---|
| **ALA** | 6.42 ± 4.28 | 4.33 ± 3.73 | 2.08 | 2.58E-03 |

| | | | | |
|---|---|---|---|---|
| **ARG** | 7.22 ± 3.91 | 4.91 ± 4.54 | 2.30 | 9.24E-04 |
| **ASN** | 7.12 ± 5.65 | 3.28 ± 2.74 | 3.84 | 3.20E-08 |
| **ASP** | 5.52 ± 3.00 | 4.69 ± 3.92 | 0.82 | 1.00E+00 |
| **CYS** | 0.03 ± 0.17 | 1.45 ± 2.02 | -1.43 | 6.92E-13 |
| **GLU** | 5.99 ± 4.64 | 9.33 ± 5.20 | -3.34 | 1.03E-05 |
| **GLN** | 5.18 ± 3.35 | 4.12 ± 3.38 | 1.05 | 3.74E-01 |
| **GLY** | 16.03 ± 6.26 | 4.87 ± 3.90 | 11.16 | 4.34E-37 |
| **HIS** | 1.01 ± 2.21 | 3.41 ± 3.02 | -2.40 | 5.64E-10 |
| **ILE** | 1.97 ± 1.93 | 4.96 ± 3.88 | -2.99 | 8.56E-12 |
| **LEU** | 3.58 ± 2.74 | 4.59 ± 4.39 | -1.01 | 6.53E-01 |
| **LYS** | 4.50 ± 3.26 | 10.78 ± 6.53 | -6.28 | 2.83E-17 |
| **MET** | 1.52 ± 1.50 | 1.20 ± 1.85 | 0.32 | 1.00E+00 |
| **PHE** | 2.97 ± 2.25 | 3.03 ± 2.96 | -0.05 | 1.00E+00 |
| **PRO** | 0.46 ± 1.12 | 1.57 ± 2.30 | -1.11 | 4.41E-05 |
| **SER** | 13.25 ± 8.53 | 6.96 ± 4.63 | 6.29 | 3.25E-09 |
| **THR** | 9.35 ± 4.73 | 12.65 ± 7.22 | -3.30 | 7.53E-04 |
| **TRP** | 0.87 ± 1.39 | 0.80 ± 1.54 | 0.07 | 1.00E+00 |
| **TYR** | 3.92 ± 2.62 | 4.69 ± 4.25 | -0.77 | 1.00E+00 |
| **VAL** | 3.11 ± 2.57 | 8.37 ± 4.48 | -5.26 | 1.58E-22 |

**Table ST7:** Amino acid composition comparison per protein region: Membrane β-barrel outward-facing vs soluble β-barrel inward-facing. Values are shown graphically in S3D.

| **Amino Acid** | **Membrane outward** | **Soluble inward** | **Difference** | **Adjusted p-value** |
|---|---|---|---|---|
| **ALA** | 11.77 ± 5.67 | 6.18 ± 4.29 | 5.59 | 2.19E-13 |
| **ARG** | 2.19 ± 2.24 | 3.62 ± 3.52 | -1.43 | 3.96E-03 |
| **ASN** | 2.07 ± 2.17 | 2.30 ± 3.21 | -0.23 | 1.00E+00 |
| **ASP** | 1.52 ± 1.63 | 1.42 ± 1.96 | 0.10 | 1.00E+00 |
| **CYS** | 0.06 ± 0.36 | 1.84 ± 2.94 | -1.78 | 1.12E-09 |
| **GLU** | 0.67 ± 1.01 | 2.82 ± 2.57 | -2.15 | 7.68E-15 |
| **GLN** | 1.88 ± 1.82 | 3.29 ± 2.96 | -1.41 | 2.54E-04 |
| **GLY** | 7.21 ± 4.88 | 3.90 ± 3.87 | 3.31 | 1.27E-06 |
| **HIS** | 1.81 ± 2.94 | 1.75 ± 2.36 | 0.05 | 1.00E+00 |
| **ILE** | 6.70 ± 4.84 | 9.85 ± 5.25 | -3.14 | 7.81E-05 |
| **LEU** | 16.05 ± 6.71 | 13.66 ± 6.96 | 2.38 | 1.77E-01 |
| **LYS** | 1.31 ± 1.55 | 2.30 ± 2.58 | -0.98 | 7.78E-03 |
| **MET** | 1.40 ± 1.77 | 4.20 ± 3.63 | -2.80 | 5.98E-12 |
| **PHE** | 8.10 ± 4.15 | 9.31 ± 4.97 | -1.21 | 8.87E-01 |
| **PRO** | 2.25 ± 3.00 | 0.63 ± 1.31 | 1.62 | 1.73E-05 |
| **SER** | 2.82 ± 2.36 | 5.17 ± 3.79 | -2.35 | 4.10E-07 |
| **THR** | 4.33 ± 3.59 | 4.93 ± 3.41 | -0.60 | 1.00E+00 |
| **TRP** | 3.84 ± 2.89 | 4.15 ± 3.96 | -0.30 | 1.00E+00 |

| | | | | |
|---|---|---|---|---|
| **TYR** | 12.06 ± 4.98 | 7.00 ± 4.06 | 5.06 | 1.54E-13 |
| **VAL** | 11.95 ± 6.84 | 11.69 ± 5.37 | 0.26 | 1.00E+00 |
